# WildCLIP: Scene and animal attribute retrieval from camera trap data with domain-adapted vision-language models

**DOI:** 10.1101/2023.12.22.572990

**Authors:** Valentin Gabeff, Marc Rußwurm, Devis Tuia, Alexander Mathis

## Abstract

Wildlife observation with camera traps has great potential for ethology and ecology, as it gathers data non-invasively in an automated way. However, camera traps produce large amounts of uncurated data, which is time-consuming to annotate. Existing methods to label these data automatically commonly use a fixed pre-defined set of distinctive classes and require many labeled examples per class to be trained. Moreover, the attributes of interest are sometimes rare and difficult to find in large data collections. Large pretrained vision-language models, such as Contrastive Language Image Pretraining (CLIP), offer great promises to facilitate the annotation process of camera-trap data. Images can be described with greater detail, the set of classes is not fixed and can be extensible on demand and pretrained models can help to retrieve rare samples. In this work, we explore the potential of CLIP to retrieve images according to environmental and ecological attributes. We create WildCLIP by fine-tuning CLIP on wildlife camera-trap images and to further increase its flexibility, we add an adapter module to better expand to novel attributes in a few-shot manner. We quantify WildCLIP’s performance and show that it can retrieve novel attributes in the Snapshot Serengeti dataset. Our findings outline new opportunities to facilitate annotation processes with complex and multi-attribute captions. The code will be made available at https://github.com/amathislab/wildclip.

## Introduction

Camera traps have become essential to monitor biodiversity (1–3) and are increasingly used for behavior research (4, 5). Camera traps are minimally invasive, also operate at night, and observe wildlife in their natural habitat. Despite these advantages, camera traps produce millions of images and remain labor-intensive to use in practice (5, 6). Traditionally, camera trap datasets are analyzed by inspecting and annotating every image according to a predefined set of attributes motivated by the scientific question of interest (Figure 1a). Depending on the study, these attributes may be the species, identities of individual animals, behaviors, or more complex phenotypical attributes. Dedicated annotation platforms are available to ease the process, but the main bottleneck remains the large quantity of data to annotate. The task gets increasingly laborious with many false triggers of the camera (due to *e.g*., vegetation movement), redundant events (*e.g*., a large herd of animals passing by), captures of small or occluded animals, or bad quality images (*e.g*., wet lens).

**Figure 1.**
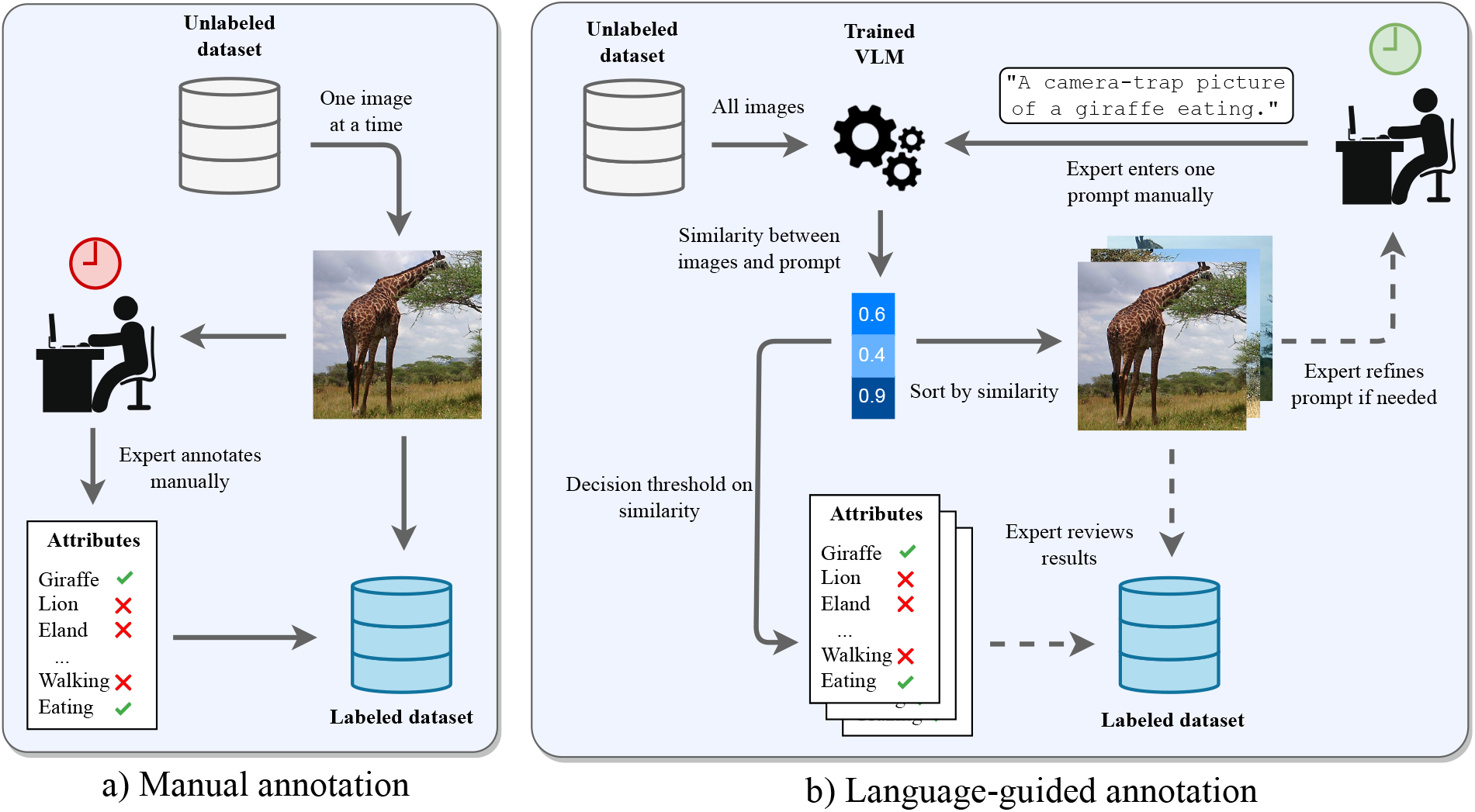
Comparison between a manual and a language-guided annotation workflow of an unlabeled camera trap dataset. a) In the manual workflow, an expert annotates sequentially every image manually. b) In the language-guided annotation workflow, an expert enters a prompt manually, which is compared to all the images in the dataset through a similarity score using a pretrained VLM. The results of such comparisons between the prompt and the images are sorted by similarity and sent to the expert to review. The expert can also iteratively refine the prompt to improve results.

To facilitate analysis, machine learning techniques can automatically filter out false positives and classify species and their behaviors (5, 7, 8). However, these classic machine learning approaches are typically trained with a predefined set of attributes (closed-set). In this work, we present a language-guided annotation pipeline that can catalyze the annotation process of an unlabeled camera trap dataset and extend machine learning analysis to potentially open sets of attributes (Figure 1b). Indeed, language naturally helps to describe events in a fine-grained fashion and facilitates the interaction between the ecologists and the model. Large Vision Language Models (VLMs) such as Contrastive Language Image Pretraining (CLIP) are particularly well suited for this task (9). Since these models were pretrained on millions of image-caption pairs, they perform remarkably well on zero-shot open-vocabulary retrieval and classification tasks (9). Yet, CLIP does not generalize well to domains substantially different from typical internet images, such as for camera trap imagery (10) or medical images (11). Consequently, several methods have been proposed to fine-tune CLIP to these specific domains (10, 12). These methods commonly fine-tune CLIP with captions that follow a fixed template and use a small vocabulary size, which inevitably degrades performance for unseen open-vocabulary captions, a phenomenon described as catastrophic forgetting (13). Ideally, the image-caption pairs used during fine-tuning should be large and diverse enough to compensate for this issue. Unfortunately, due to the temporal burden in annotating datasets, camera-trap images are rarely labeled beyond species-level annotations (14–18) and the set of possible labels to construct image captions from remains constrained to a small vocabulary. One notable exception is the Snapshot Serengeti dataset that benefited from a citizen science initiative that provided more detailed information for each image (19).

In this work, we present an adaptive framework for CLIP to the domain of camera trap images (WildCLIP) that we evaluate on Snapshot Serengeti (19). To mitigate the problem of catastrophic forgetting, we follow a recently proposed vocabulary-replay method (20). Based on automated literature search, we create a replay vocabulary relevant to the domain of interest and use it to preserve the structure of the embedding space during training. We also build upon CLIP-Adapter (12) to dynamically add new vocabulary to the model with few labeled samples. The open-vocabulary performance of the method is quantitatively evaluated on held-out words and caption templates. We also provide qualitative results for open-set queries inspired by what an ecologist might use. We also explore how our training strategy allows the model to dissociate between the species and their context. Specifically, our contributions are the following:

- We create WildCLIP by fine-tuning CLIP to retrieve images corresponding to diverse attributes and environmental conditions from camera-trap datasets and benchmark it on Snapshot Serengeti.
- Through a series of quantitative and qualitative examples, we analyse the behavior of WildCLIP in details, also focusing on zero- and few-shot abilities on open vocabulary.

We hope that our work motivates the creation of richly annotated camera trap datasets, to collectively create powerful VLMs for camera trap data.

## Background and related works

### Machine learning applications for camera trap imagery

Applications of machine learning to camera trap data mainly focused on animal tracking (21, 22) and species recognition (23, 24). During the last decade, the development of convolutional neural networks (CNNs) largely improved the performance of vision models for animal detection (7, 22, 25, 26), species classification (17, 27–30), behavior recognition (8, 31) or animal counting (8, 32). In 2018, Norouzzadeh et al. (8) showed an innovative pipeline to classify species, count animals and assess age, behavior, and interactions with other individuals from the Snapshot Serengeti consensus data, still making it one of the most diverse multilabel classification method for camera traps to date.

However, other tasks such as assessing animal body conditions, have received less methodological focus from the deep learning community, despite the interest from ecologists (33–35). This absence of research is partly attributable to the lack of publicly available annotations beyond taxonomies. This is related to the difficulty in crowd-sourcing such attributes, as they can be subjective, undergo subtle variations and may require substantial expertise. In these cases, active learning is a way to compensate for the lack of labels (36–39). However, this approach requires a few annotated samples to initiate the process, which may be difficult to find for rare events. Few-shot learning and self-supervised learning also promise to improve the data efficiency (40). A more recent way to learn in low-label regimes is to use VLMs pretrained on millions of image-text pairs.

### Large scale multi-modal language models

With the advent of transformers (41), large language models (LLMs) emerged that demonstrated remarkable capabilities for natural language processing tasks incl. ChatGPT (42–45). LLMs can also be used to exploit pre-trained AI models to carry out various tasks (46, 47) including behavioral analysis (48). Concurrently, multi-modal variants were also created, in particular large scale visual-language models, which have tremendously improved the performance and robustness for zero-shot object recognition, image search and many other tasks (9, 11, 49–51). One of the earliest models in this domain was CLIP (9), which can be tuned to related domains of interest with CLIP-Adapter (12). Here, a Multi Layer Perceptron (MLP) modulates the vision feature vectors and is added at the end of the vision backbone and weighted by a parameter *α*. The method is then trained with a cross-entropy loss. Similarly, Pantazis *et al*. (10) proposed the Self-supervised Vision-Language Adapter (SVL-Adapter) and demonstrated that fine-tuning is needed to adapt CLIP to the domain of camera traps and presented a method with improved performance over CLIP-Adapter for few-shot species classification on challenging camera trap datasets. Their method blends the class probabilities of CLIP with the output of an additional vision backbone trained with self-supervised learning. This has the disadvantage of limiting the method to a fixed set of queries during training and at inference, here corresponding to the set of species.

As mentioned in the Introduction, fine-tuning CLIP with a small vocabulary size will inevitably limit its use for open vocabulary queries. To mitigate this issue, Ding *et al*. proposed a vocabulary replay method abbreviated as VR-LwF to prevent the model from forgetting concepts related to a task of interest (20). The method stems from the “Learning without Forgetting” (LwF) approach to catastrophic forgetting (52), and exploits the alignment between text and image modalities of CLIP to circumvent the need for annotated image-caption pairs. Specifically, a loss term is added during training that minimizes the distribution shift of the cosine similarities between training image embeddings and the text embeddings of an arbitrary set of words referred to as “Vocabulary Replay” (VR).

### Background on CLIP

Contrastive Language-Image Pretraining (CLIP) is a VLM for open-vocabulary classification tasks (9). It consists of a visual encoder (VE) and text encoder (TE). The similarity metric for image **x**_*i*_ and caption **y**_*j*_ is computed as:

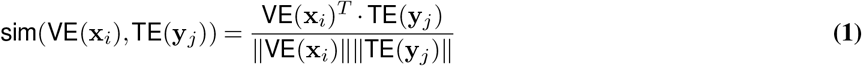

CLIP was trained to learn a joint embedding space for image and text representations using a contrastive loss on millions of image-caption pairs (9). During training, each batch of size *N* ^2^ is composed of *N* positive image-caption pairs, and the remaining *N* × (*N* − 1) are considered negative pairs. The loss aims at maximizing the similarity of the positive pairs and minimizing it for negative pairs:

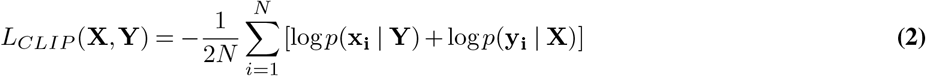

Here, the likelihoods following Equations (3 and 4), where *τ* is the temperature parameter:

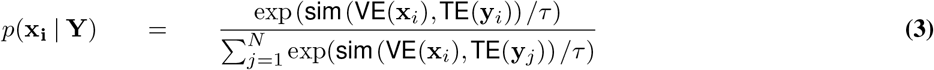

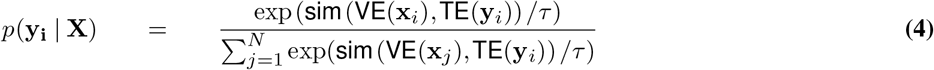

At inference time, CLIP is used to compute the cosine similarity between queries and images. If queries correspond to mutually exclusive classes (*e.g*., “A camera trap picture of a `<class_name>`“), a softmax operation is commonly applied to return respective class probabilities.

## Methods: WildCLIP and WildCLIP-Adapter

Our method consists of two steps: first, we fine-tune the vision encoder of CLIP on a large dataset of camera trap images and their associated captions (Figure 2a). Second, we freeze the vision encoder and train a Multi-Layer Perceptron (the “Adapter (12)) with a few samples of sequence-caption pairs to learn words from a *novel vocabulary* (Figure 2b). In other words, the first step fine-tunes CLIP to a WildCLIP model with a more fine-grained representation of camera-trap imagery using a closed-set domain of common queries from a base vocabulary (Figure 2a). The second step adapts WildCLIP towards an open set of queries that a trained domain expert can provide interactively. To further preserve open-vocabulary capabilities of CLIP, we add an extra loss term (20) that replays vocabulary related to the domain of interest (Figure 2b). Eventually, our method allows its users to dynamically query and explore camera trap imagery (Figure 2c).

**Figure 2.**
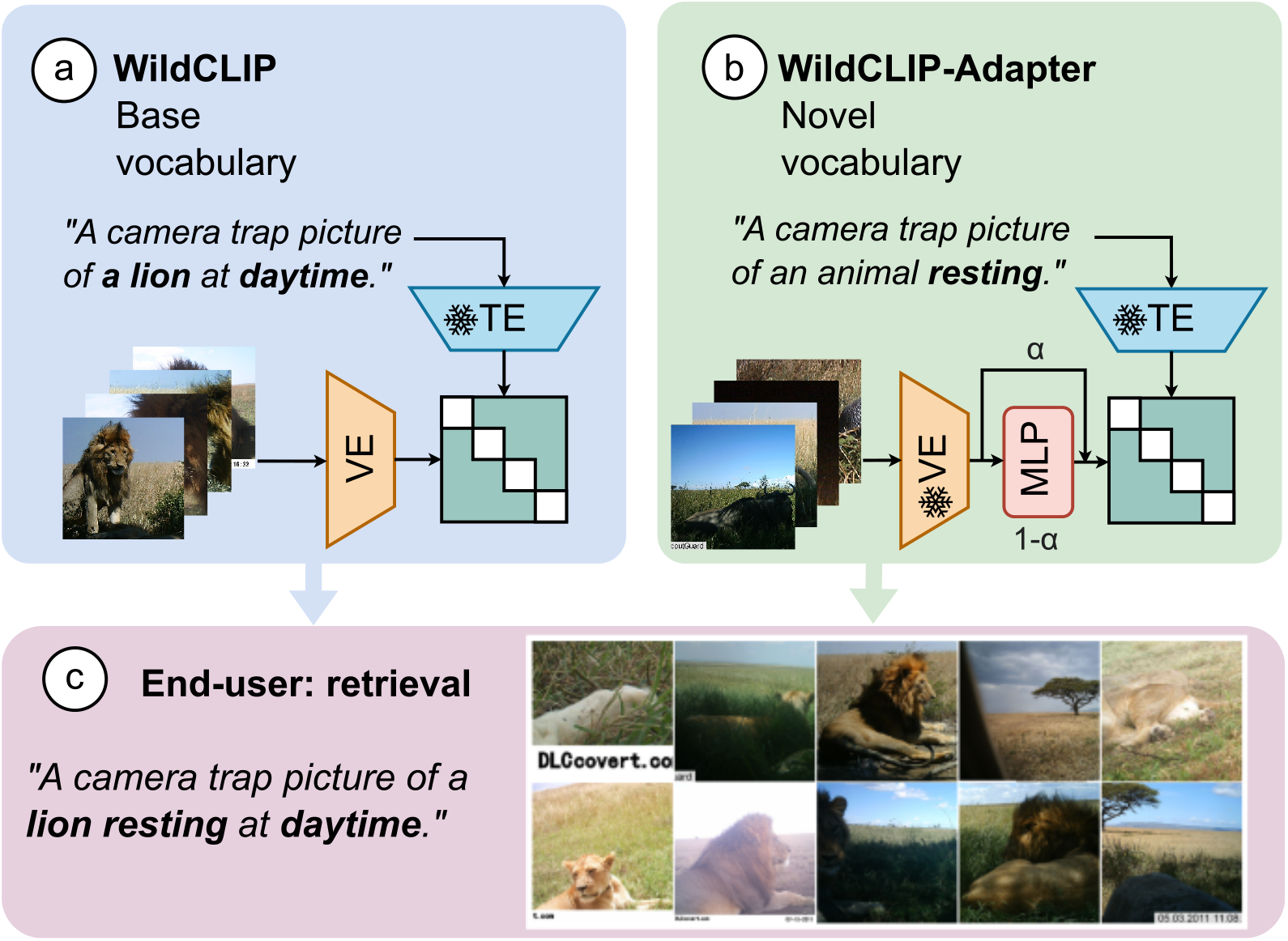
WildCLIP and WildCLIP-Adapter. We fine-tune CLIP to the domain of camera-trap datasets by fine-tuning its visual encoder with augmented image-caption pairs (a). We further adapt the model with an MLP adapter on a novel set of words to demonstrate the advantage of using VLMs (b). Finally, we evaluate how these two models can be used for image retrieval on a set of novel images (c).

### Fine-tuning (WildCLIP)

We use CLIP’s original contrastive loss (Equation 2) to fine-tune the CLIP-pretrained visual backbone (9). The text encoder is kept frozen to avoid forgetting the open-vocabulary knowledge of CLIP. We create multiple captions for every image using multiple caption templates and the available image labels. Specifically, we generate all possible combinations of labels describing an image, and apply them to ten different caption templates (see Figure 4 for examples). This process significantly depends on the available labels and is further discussed in the “Experimental set-up” below. We use up to seven caption templates for training, and leave the remaining ones for evaluation. We hypothesize that training on multiple templates will make the model robust to different formulations of queries. On the other hand, a model trained with only one template may overfit (on this one).

The set of augmented image-caption pairs becomes inevitably unbalanced if some labels describe multiple images, which adds to the natural imbalance of camera trap datasets. We balance the dataset of image-caption pairs with a mix of upsampling and downsampling so that rare captions appear as often as common ones. We use data augmentation on the colors and the geometry of the image to increase visual diversity, which has been shown to improve generalization.

### Few-shot adaptation (WildCLIP-Adapter)

In this step, we expand the WildCLIP vocabulary to new words, following a similar approach as Gao et al. (12). We add a two-layer perceptron with a residual connection, weighted by a fixed parameter *α*, at the end of the pretrained visual encoder of WildCLIP. This perceptron adapts the image representation vectors to the new vocabulary so that they better align to the frozen text vectors of WildCLIP, while still keeping information from the base vocabulary. Differently from (12), we input image-text pairs to the model, and we use a custom loss that maximizes the cosine similarity between the positive pairs only (*i.e*., the diagonal elements of the text-image features alignment matrix). This is motivated by the observations that captions can have multiple matching images and *vice versa*, yielding several false negative pairs in every batch, which is a problem for few-shot learning. As we expect performance to be sensitive to the choice of the few-shot samples used for adaptation, we repeat the experiment 5 times with different image samples from the novel vocabulary set and report the mean in the results. We refer to our modified version of CLIP-Adapter as CLIP-Adapter∗.

### Addressing catastrophic forgetting (VR-LwF)

As discussed in Related Works, fine-tuning CLIP on a fixed vocabulary may reduce its open-vocabulary abilities. When fine-tuning CLIP with the vocabulary of WildCLIP, we can view the embedding space as shrinking towards the volume containing training caption embeddings only (Figure 3a). Even though we do not fine-tune the CLIP text encoder (TE), the vision encoder (VE) will only learn to match images with a small set of captions. This shrinking is responsible for the catastrophic forgetting. On the contrary, we aim at expanding the latent space learned by WildCLIP also to contain vocabulary relevant to the task of interest, here ecology, denoted as CLIP_*ecology*_, while still forgetting totally irrelevant concepts. To achieve this, we follow the VR-LwF method of (20). Specifically, we replay relevant vocabulary through the TE, that we refer to as text “anchors”, since the text encoder is kept frozen. Since the pool of anchors **A** is noisy, some fall outside of CLIP_*ecology*_, while others are already contained within WildCLIP’s vocabulary. We then ensure that the distance between the images and the anchors does not drift too much in the latent space during training (Figure 3b).

**Figure 3.**
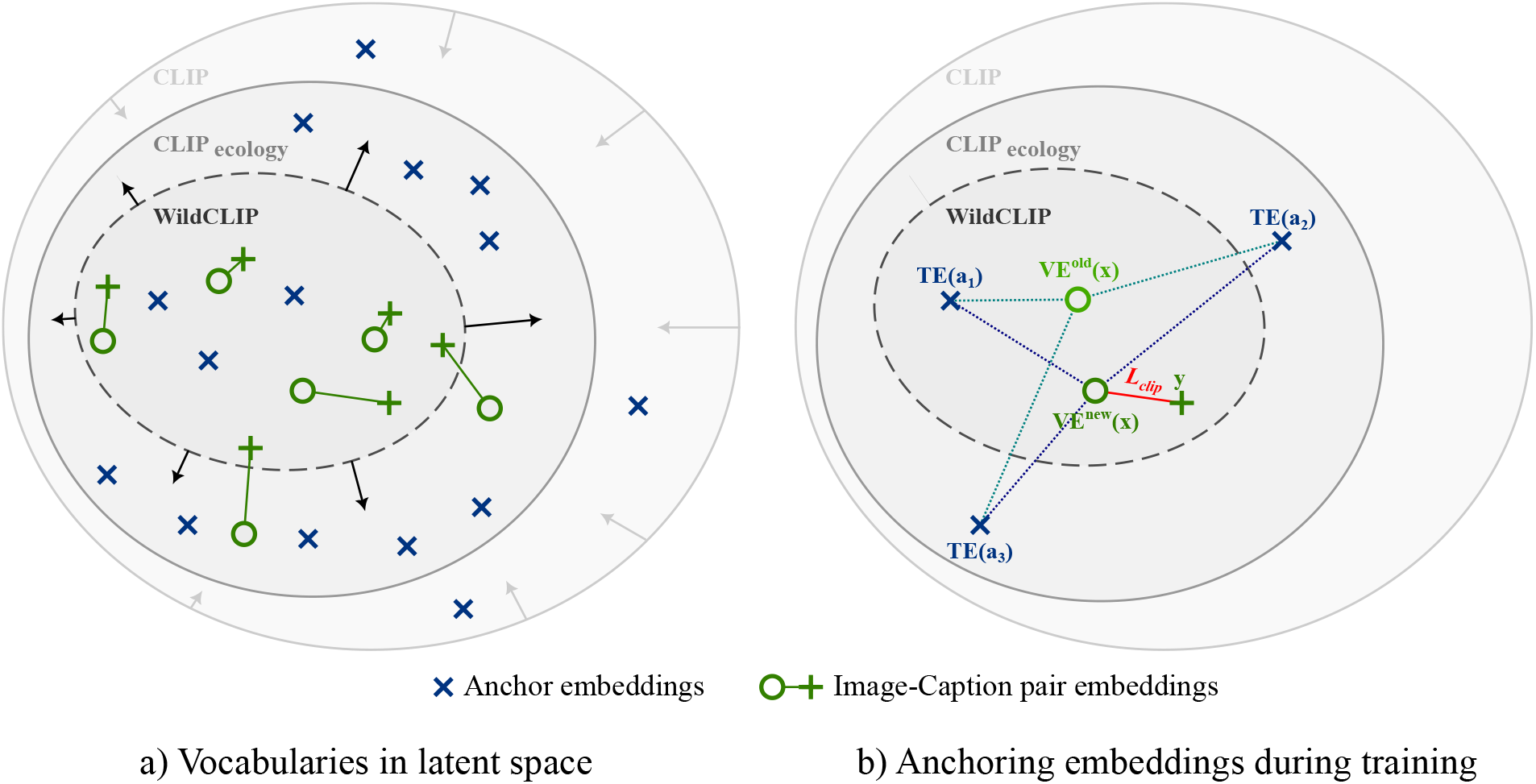
Embedding space when applying VR-LwF to WildCLIP for a given image-caption pair. a) CLIP embedding space contains many different concepts unrelated to our task. We aim at using vocabulary replay to learn embeddings in the domain of interest (CLIP_ecology_) while only having captions embeddings in the WildCLIP embedding space. b) For a given image-caption pair **x** − **y**, we compute the cosine similarities of the previous VE^old^(**x**) and new VE^new^(**x**) image embeddings with respect to all replayed vocabulary embeddings TE(**a**_*j*_). We also compute the usual cosine similarity *L*_*CLIP*_ (**x, y**) (Equation 2) between the new image embedding and the matching caption text embedding. By minimizing the cross-entropy between cosine similarity distributions, we expect the VR-LwF method to preserve some open-vocabulary capabilities of CLIP. This loss term is counter-balanced by *L*_*CLIP*_, which aims to minimize the cosine similarity between positive image-caption embeddings.

In practice, for each image **x**_*i*_ of a given batch of *N* positive image-caption pairs, we compute the distribution of cosine similarities of **x**_*i*_ embeddings with respect to the pool of anchors **A** of size *N*_*A*_ when **x**_*i*_ is passed through the previous vision encoder (VE^*old*^) and the one being trained (VE^*new*^), denoted as 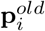 and 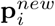, respectively (Figure 3b, dotted lines). We then compute the 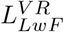 loss as the cross-entropy between both distributions and minimize its sum over all images:

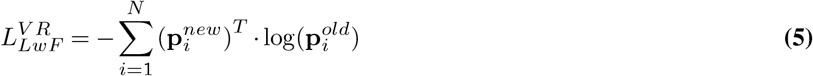

with probabilities:

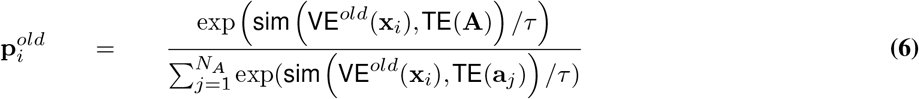

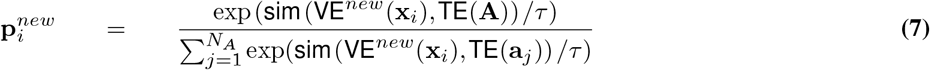

The final training loss is the sum of *L*_*CLIP*_ (Equation 2) and 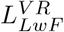 (Eq. 5).

## Experimental set-up

### Data

The Snapshot Serengeti camera-trap dataset (19) was collected over eleven seasons since 2010 and contains more than seven million images from the Serengeti national park, Tanzania. The dataset benefited from large-scale annotations from a citizen science initiative.

#### Species labels

We use MegaDetector (7) outputs from seasons 1-6 provided on LILA BC. We restrict our study to sequences containing single individuals only since consensus multilabels are provided at the sequence level without distinctions between individuals.

#### Behavior labels

Behavior labels are reported as the proportion of users who voted for a given behavior. We set the behavior visible in an image as the behavior with the most votes. Since we consider single individuals only, the “*Interacting*” behavior is removed. We set the age label to “*Young*” if more than 50% of the users voted for the category “*Baby* “.

#### Scene labels

Because the Serengeti Park is relatively close to the equator, we label images taken between 6 a.m. and 7 p.m. as “*daytime*” and as “*nighttime*” otherwise, independently of the month. For the camera environment, we manually annotated whether a camera field of view is pointing towards “*grassland* “ or “*woodland* “.

In the end, each sample image is described by five attributes: 1) the depicted species, 2) its age, 3) its behavior, 4) a binary day/nighttime label, and 5) the environment surrounding the camera (“*grassland* “ or “*woodland* “). Further details on image pre-processing are detailed in the Appendix.

### Building image captions and test queries

From the five attributes describing each image, we automatically build structured captions following ten different templates (Figure 4, see Appendix). Given a set of attributes, corresponding captions built from different templates all express the same information but with a different formulation (*e.g*., ordering of the attributes in the sentence or contextual words.) We create every possible and unique combination of captions with respect to the attributes and the different templates, yielding 297 captions per image.

**Figure 4.**
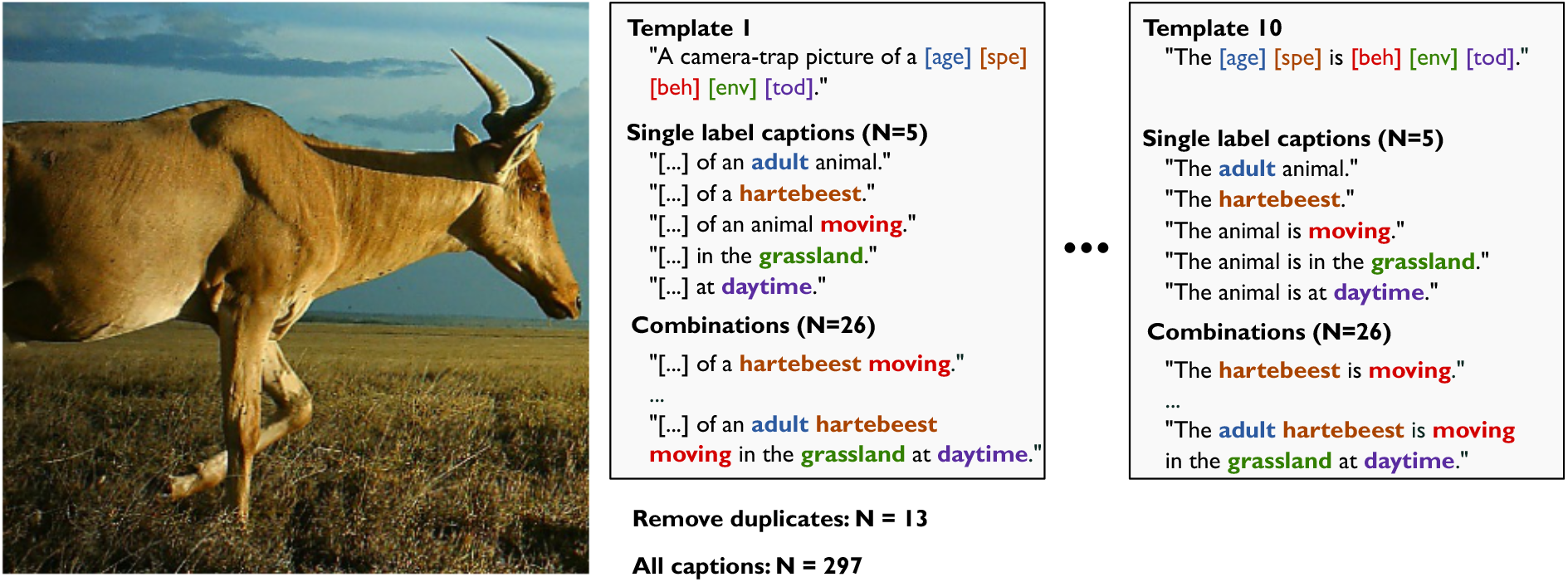
Building image captions. 297 structured captions following 10 different templates describe each image.

### Replay Vocabulary

We build an external set of words relevant to the Serengeti wildlife to preserve the representation of concepts not associated to an image during fine-tuning. To do so, we automatically parse the title of ecology papers related to Serengeti wildlife and extract keywords. Following (20), we build 100 5-grams composed of these keywords by randomly sampling them without replacement. These 5-grams constitute the pool of anchors **A** introduced in Methods. More information on the creation of the replayed vocabulary and examples of 5-grams can be found in the Appendix.

We note that the retrieved vocabulary extends beyond the domain of interest, with vocabulary including politics and virology (Appendix). Although unrelated words and random sentences may seem inefficient, we assume that VR-LwF is robust to the chosen anchors (see Results). Since this method prevents the model from overfitting to the vocabulary, it is fine-tuned on by constraining the drift of the vector embeddings in the latent space, we hypothesize that the choice of the words matters less than their embeddings evenly spanning a volume of the latent space that relates to the task of interest (See Figure 3a, CLIP_*ecology*_).

### Data split

We divided images into training and testing partitions, as well as the split of the captions into two sets of vocabularies (Figure 5). Training and testing images are split at the camera level following recommendations from LILA (53). WildCLIP is trained with samples from the *base vocabulary*. This set contains images of species like *“Thomson’s gazelle”, “topi”*, or *“ostrich”* in different scene and behavior settings like *“daytime*”/*”nighttime*”, *“eating*”/*”moving*”. WildCLIP-Adapter is then further trained with up to 8 sequences of 1 to 3 images for each caption from the *novel vocabulary*. Crucially, the novel vocabulary contains different species like *“Grant’s gazelle*”, *“leopard* “, behaviors like *“standing*” and *“resting*”, and the two different habitats *“woodland* “ and *“grassland* “. To preserve independence, we ensure image-caption pairs containing the novel words are never seen during the training of WildCLIP.

**Figure 5.**
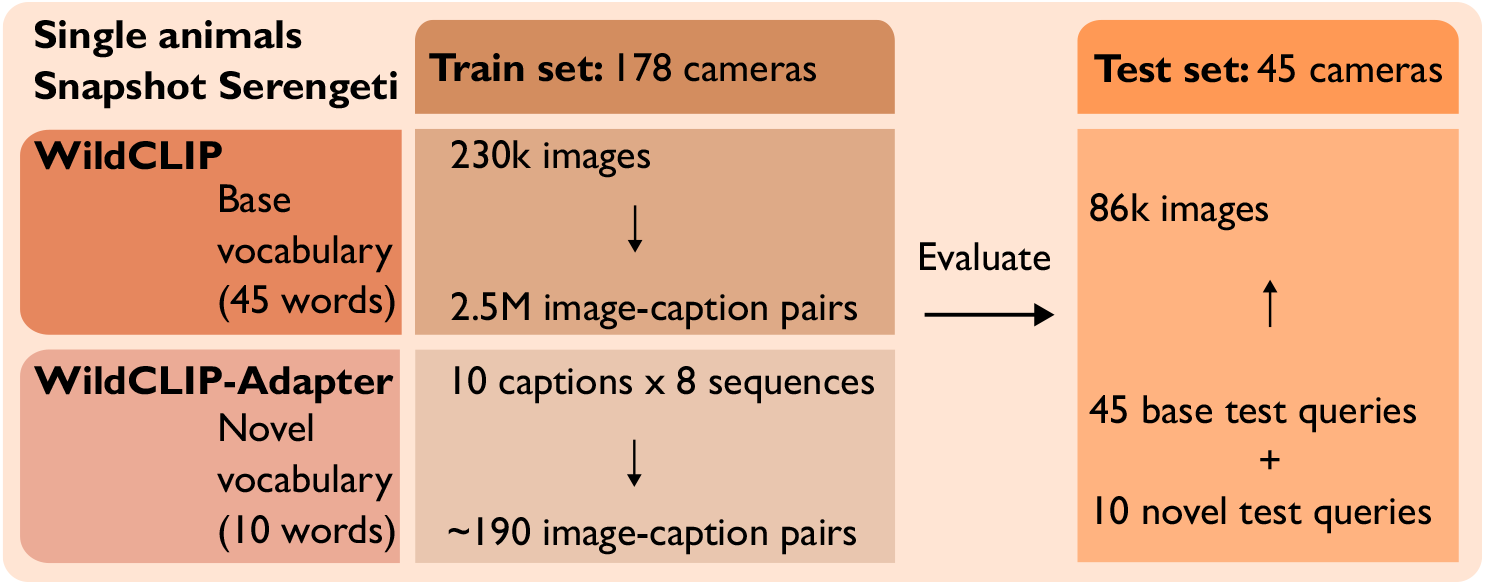
Data split for quantitative evaluation. WildCLIP is trained on base vocabulary (top left) and adapted further to WildCLIP-Adapter with novel vocabulary captions (bottom left). Test data is split on a camera level, and both models are evaluated on base and novel vocabulary separately on images of 45 new camera traps (right). The number of training image-caption pairs and of test queries is computed according to template 1 only.

We also split the test queries into “in-domain” templates and “out-of-domain” ones. WildCLIP is trained on either only on template 1 (*t*_1_), or on templates 1 to 7 (*t*_1−7_), and its performance is evaluated on either “in-domain” template 1, or on “out-of-domain” templates 8 to 10 (*t*_8−10_).

### Evaluation metrics

We evaluate WildCLIP as a retrieval task, meaning that for a given test query, the true corresponding images should rank higher in cosine similarity with the test query than non-matching images. The set of test queries for the retrieval task is defined as the set of structured captions containing single attributes, yielding a direct equivalence between individual multilabels and test queries, for which performance can be measured. Note that WildCLIP is not limited to these single attribute captions, as it can retrieve images at every level of complexity (which is the method’s main advantage); nevertheless, here, we limit our test captions to single attributes to allow direct comparisons to finetuned models. We compute the mean average precision (mAP) from the alignment scores per test query and then average over all test queries.

### Ablation study

We control the performance of the different additions to our method with an ablation study, considering CLIP ViT/B-16 performance as our baseline.

To evaluate the effect of adding language when learning the representation of camera trap images, we first compare WildCLIP with the pretrained visual backbone of CLIP, to which an MLP head has been added with binary output neurons corresponding to each possible test query from the base set (ViT-B-16-base). We also report the performance of this model on the novel vocabulary (ViT-B-16-novel) by replacing the output layer of the pre-trained model with an output layer with 10 output units (fixed size of the novel vocabulary in this setup) and adapting it with the same few-shot scenario as for WildCLIP-Adapter∗ and CLIP-Adapter∗, but using a binary cross-entropy loss.

To further motivate our approach over existing ones, we train CLIP-Adapter∗ (see Methods), where only the additional MLP head is trained, and the backbone of CLIP is kept frozen. Since training a vision transformer is computationally expensive, we evaluate the choice of the visual backbone by comparing performance between a ResNet50 backbone with the default ViT/B-16 one.

To assess the generalization to out-of-domain template structures (templates 8 to 10, see Methods) for the test queries, we compare the performance of WildCLIP when trained on a single (template 1) or on seven templates (templates 1 to 7). Finally, we assess the effect of the VR-LwF loss during fine-tuning and during adaptation.

## Results

We start by showing qualitative results of WildCLIP, contrasting it with CLIP. Then we will evaluate the performance and carry out an ablation analysis.

### Qualitative results for complex queries

We illustrate how WildCLIP improves on CLIP when retrieving images using complex queries which have been seen during training (Figure 6). Looking at the retrieval results one can note that CLIP already performs well for queries containing only the species name (*e.g*., *“a giraffe”*), but sometimes fails when additionally prompted with behavioral information (*e.g*., *“a giraffe eating”*). On the contrary, WildCLIP generally performs well for these complex queries. For the novel query *“A camera-trap picture of a male lion resting at daytime.”*, WildCLIP-LwF-adapter-LwF best retrieves the corresponding events, where *“resting”* is a word from the novel vocabulary. Despite the VR-LwF loss, this still comes with a decreased retrieval performance on queries from the base vocabulary such as *“A camera-trap picture of a giraffe.”* More qualitative examples can be found in Appendix Figure 10.

**Figure 6.**
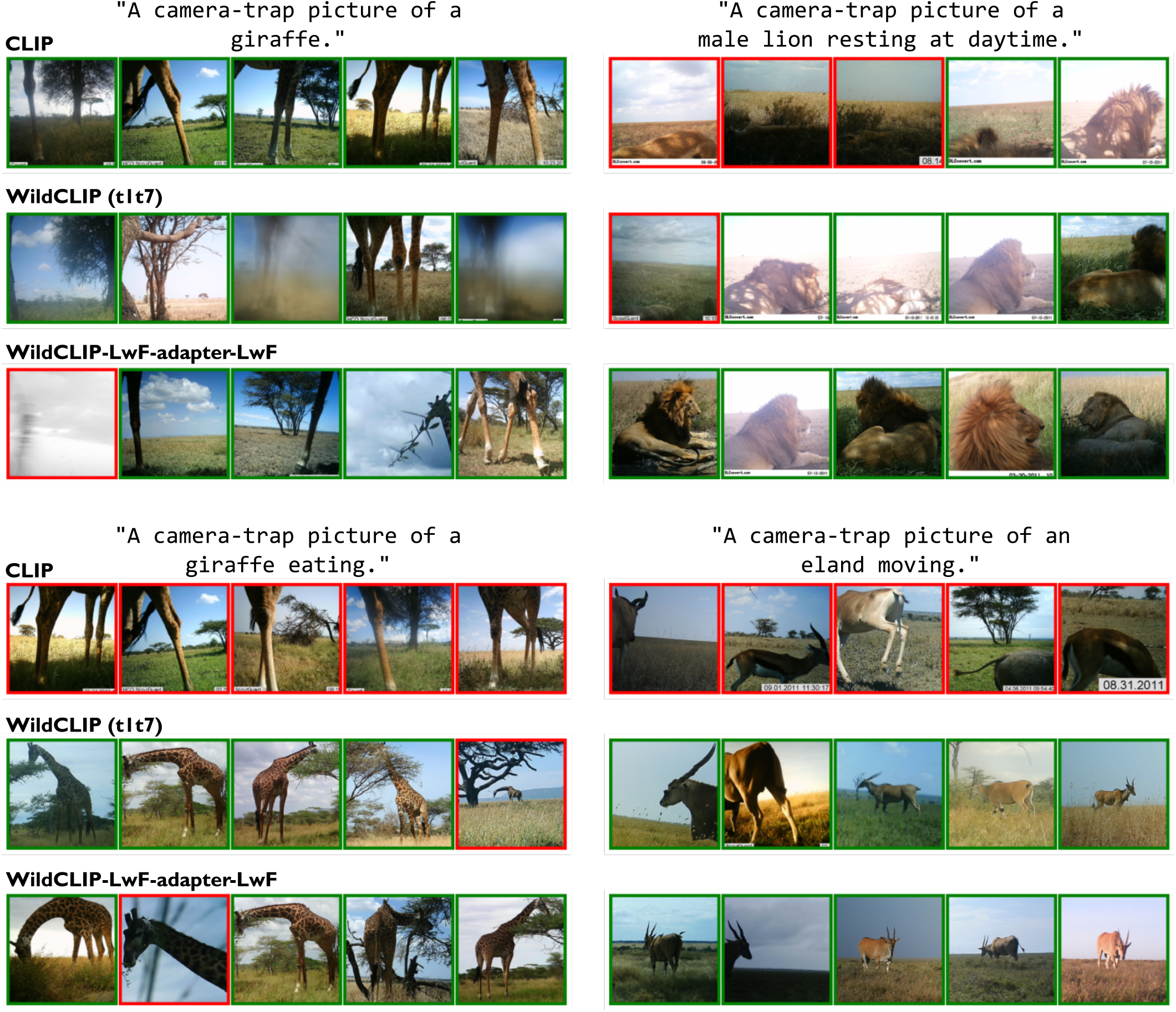
Qualitative results on complex queries. Top-5 test images most aligned with the given complex queries. *”Resting”* is a word from the novel vocabulary.

Having different captions describing a single image may seem misleading for the model. However, we hypothesize that it helps the model disentangle the multiple attributes of this image. Indeed, for WildCLIP, the top-3 captions most similar to the waterbuck images are a combination of species, behavioral, and environmental information (Figure 7). In contrast, CLIP only retrieves species information. This suggests that CLIP mainly learned to associate captions describing an object from an image, disregarding contextual information. We explored this disentanglement further. We progressively modify the input query by modulating contextual or behavioral information. We observe coherent changes while the species retrieved remains unchanged (Figure 8). This qualitatively suggests that our method successfully retrieves events with a detailed level of contextualization. We see that the model reaches its limit for the grassland environment, which is part of the novel vocabulary on which WildCLIP-LwF was not fine-tuned. Even though the animals are in the grassland, they are not all topis, and two are not eating.

**Figure 7.**
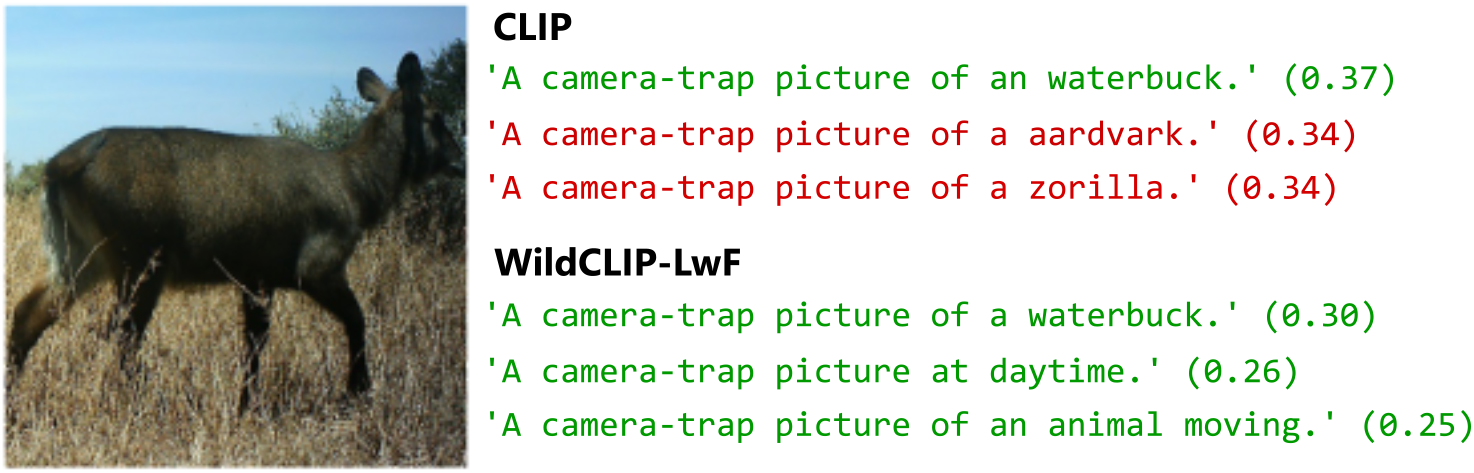
Top-3 test queries most aligned with the image for WildCLIP along with alignment similarities

**Figure 8.**
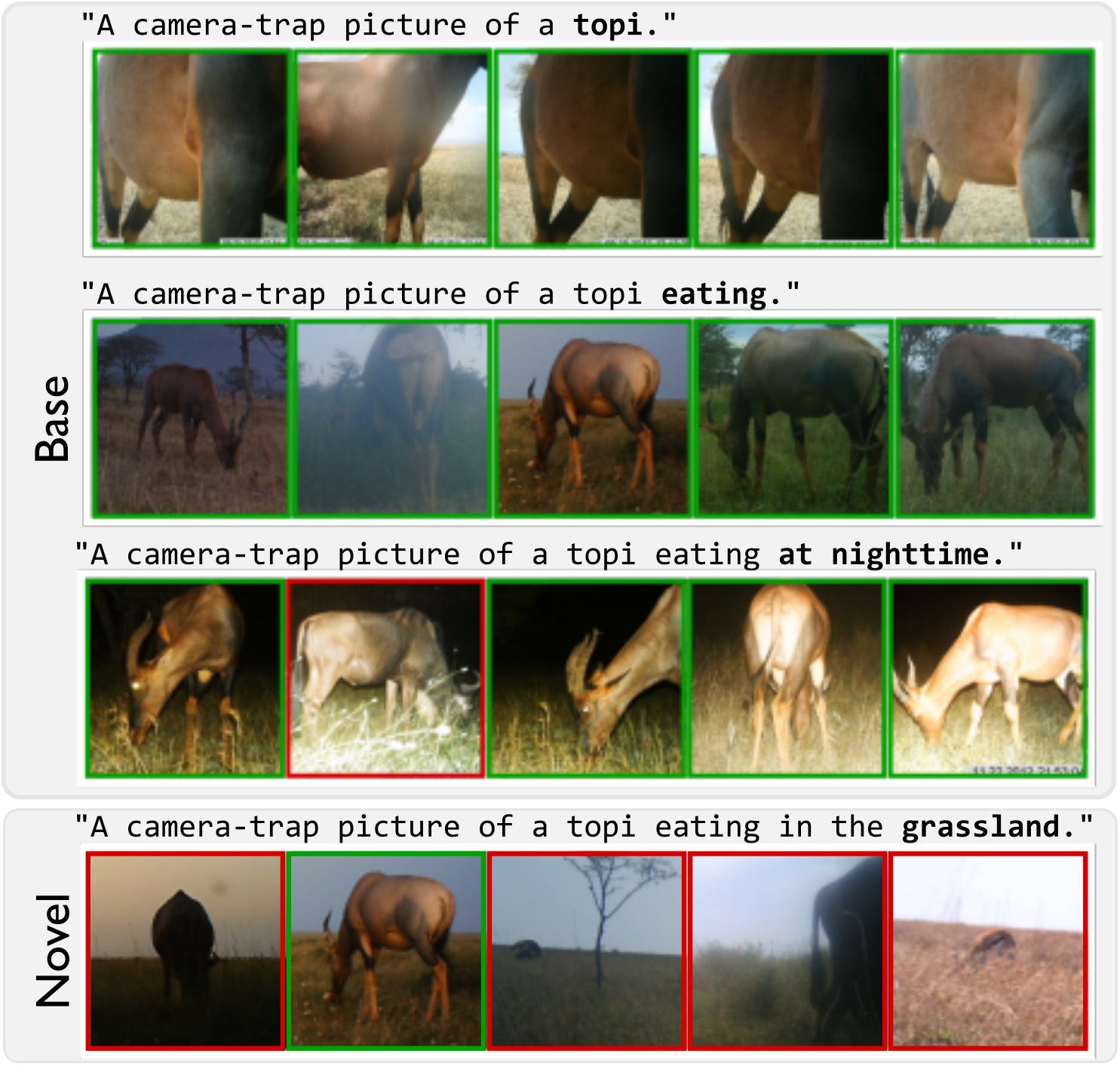
Top-5 most similar images for WildCLIP-LwF to complex queries by progressively adding or modifying some attributes from the base and the novel vocabularies (bold).

### Open-vocabulary qualitative results

Qualitative results illustrate the potential of WildCLIP to retrieve events of interest from open-vocabulary queries (Figure 9). Here we compare the original CLIP retrieval performance with WildCLIP pretrained on seven templates and the same model further trained on 2 to 8 shots of the proposed captions (only two samples of hyena with a carcass were observed in the subset of the train set, see Methods). We observe a clear qualitative improvement from CLIP to WildCLIP for the prompt: *“A hyena carrying a carcass.”*, with 4 retrieved events within the top-5, and 4 for WildCLIP-Adapter-LwF as opposed to one visible carcass for CLIP. WildCLIP also performs better on the attribute “dry grass”. However, the original CLIP qualitatively outperforms the trained model for the running behavior and the animal’s position on the camera. These results suggest that when CLIP already retrieves corresponding events for unseen open-vocabulary queries, WildCLIP do not improve much or may even reduce performance. On the other hand, we see improvements in cases where CLIP fails. This further motivates us to improve the proposed methods to preserve the original embedding space (VR-LwF (20)) and to retain some of the original CLIP embeddings (CLIP-Adapter (12)). We also provide more zero-shot qualitative examples for CLIP, WildCLIP and WildCLIP-LwF in Appendix Figure 11.

**Figure 9.**
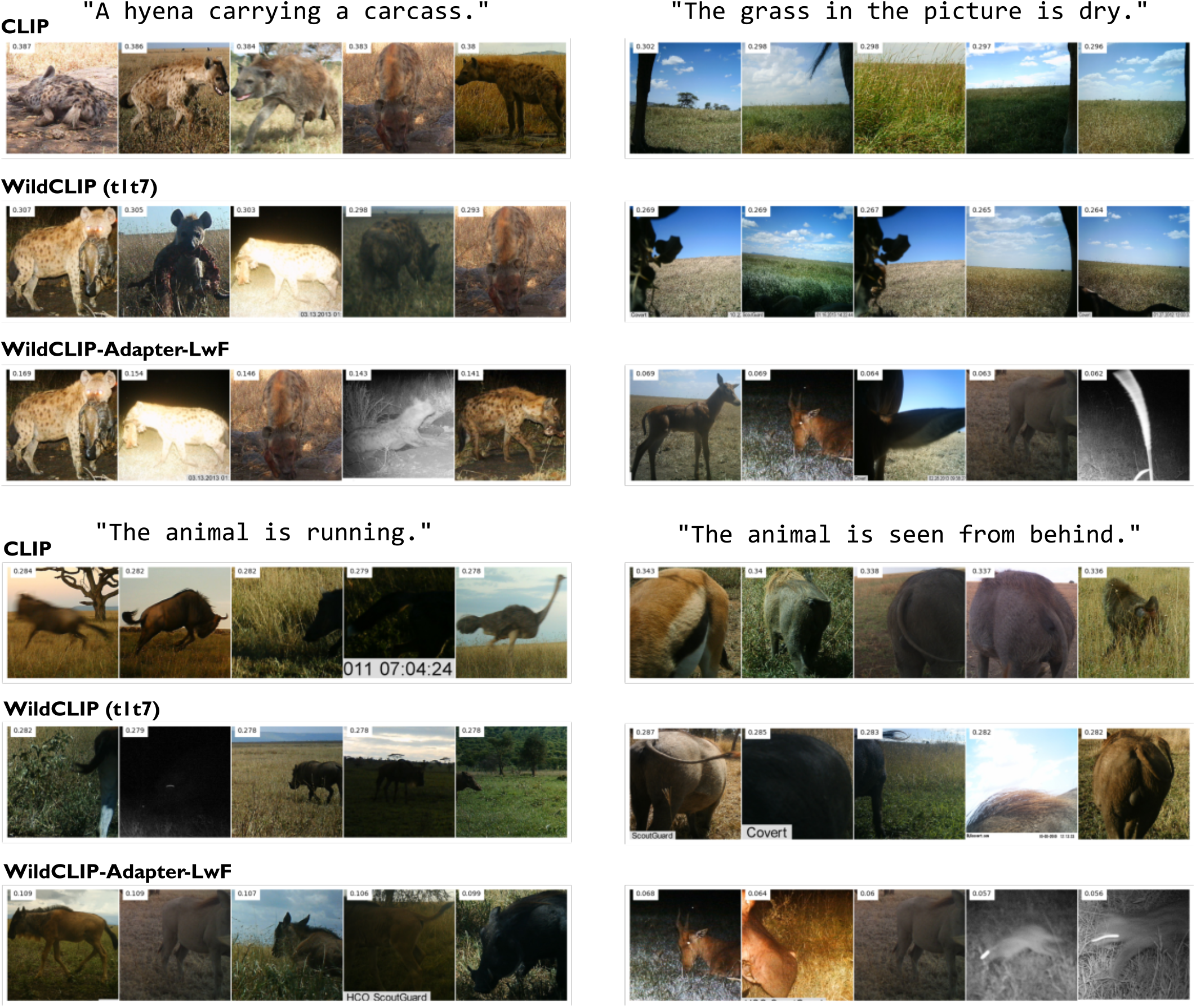
Qualitative results on open vocabulary queries. Top 5 most similar images to each given query. For each query; *first row* : Original CLIP model; *second row* WildCLIP pretrained on templates 1 to 7; *third row* WildCLIP is further trained following the WildCLIP-Adapter∗ methodology (see Methods) on 2 shots (top-left) and 8 shots (others) of these queries.

After illustrating promising capabilities of WildCLIP as well as failure cases, we sought to rigorously evaluate its performance.

### Quantitative Comparison

Our full method, WildCLIP-LwF, significantly outperforms CLIP on the image retrieval tasks (Table 1), showing that the model is better adapted to the domain of camera traps. Indeed, we see an improvement of +0.31 for WildCLIP-LwF over CLIP for the base vocabulary. Importantly, fine-tuning also improves the performance on the novel vocabulary (+0.12), although WildCLIP-LwF was not trained on these words. WildCLIP-LwF-Adapter∗-LwF does not improve on WildCLIP-LwF for the novel vocabulary, but still improves on CLIP by +0.08.

**Table 1.**
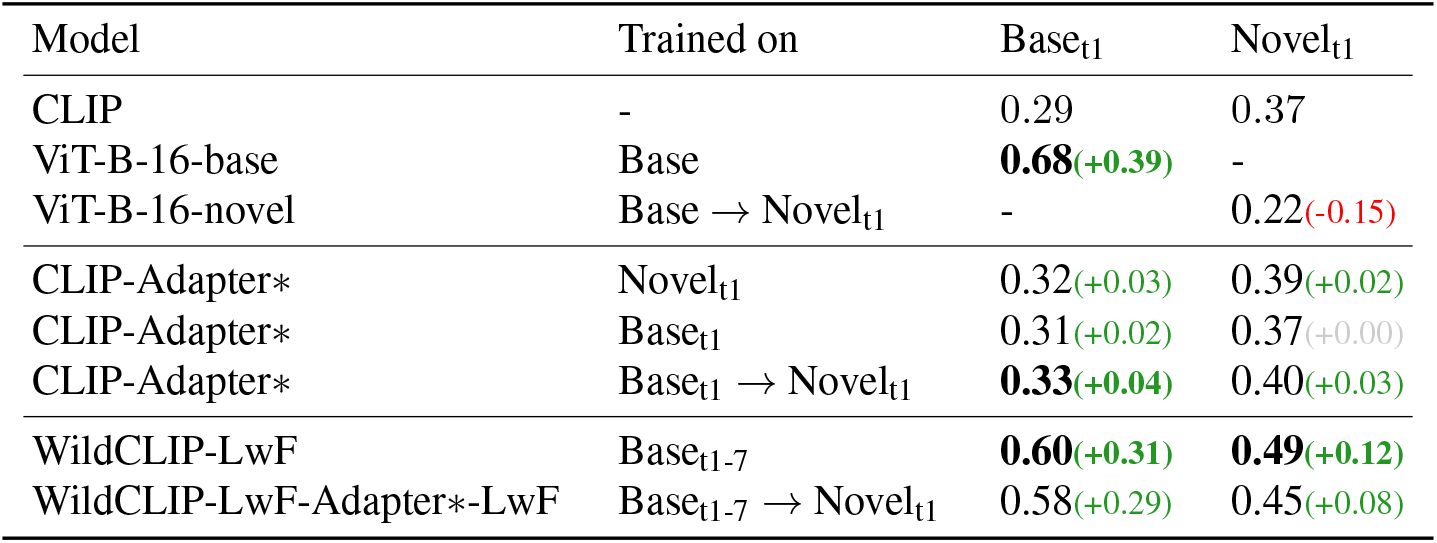
Mean average precision (mAP) and difference from CLIP on base and novel vocabularies of the test set. The performances of models trained on the novel vocabulary are reported as the average of the five repetitions of the 8-shots training, but standard deviation is not repeated for readibility and was consistently below 0.01. Clip-Adapter∗ is adapted from (12) as explained in the Methods. Arrows (→) denote adaptation. Dataset subscripts denote used captions and queries templates as described in the Methods. Models are all pretrained with CLIP ViT-B/16.

We also compare WildCLIP to CLIP-Adapter. We see a significant advantage of fine-tuning the entire visual backbone of CLIP (WildCLIP-LwF, Table 1) over learning a new MLP head only (CLIP-Adapter∗), when training them both on the base vocabulary. WildCLIP-LwF-Adapter∗-LwF also performs better than CLIP-Adapter∗ on both the base and the novel vocabularies after 8 shots (+0.29 *vs*. +0.02). This corroborates the results from Pantazis et al. (10) that CLIP should be adapted for camera trap data. Furthermore, our method significantly outperforms CLIP-Adapter∗.

Finally, we also compare to vision-only models in the classic transfer learning setting. The performance of a vision-only model pretrained from the CLIP visual backbone is slightly above the performance of WildCLIP-LwF on the base vocabulary (0.68 *vs*. 0.60). This is most likely due to the different loss functions (contrastive loss and binary cross entropy, respectively), where a vision-only model is not constrained to match the learnt image embeddings to frozen text ones. However, the performance of WildCLIP-LwF-Adapter∗-LwF surpasses the one of the vision model (0.45 *vs*. 0.22). Overall, this suggests that using a VLM for the retrieval task instead of a closed-set, vision-only model slightly decreases the performance, while providing all the advantages of dynamically interacting with the dataset through text, including easy and accurate adaptation to new vocabularies, while the vision-only model cannot.

### Ablation study

We carried out a number of ablations to justify our design decisions. Firstly, we will ablate different components of WildCLIP-LwF.

#### Backbone

Firstly, for the original CLIP model, a vision-transformer backbone improves the ResNet50 backbone performance by around +0.05 on both base and novel vocabularies (Table 2). This is consistent with results reported in Radford et al. (9). A consistent result is also observed when training WildCLIP, although the performance boost is mainly visible on the out-of-domain test query templates for both base and novel vocabularies.

**Table 2.** Ablation Study. We test the effect of different visual encoding backbones, the impact of the trained templates and the mutual effects of the VR-LwF loss (20) and the adapter module (12). Unless stated otherwise, CLIP visual backbone is the pretrained ViT-B/16 one.

| Model | Trained on | Base<sub>t1</sub> | Novel<sub>t1</sub> | Base<sub>t8-10</sub> | Novel<sub>t8-10</sub> |
| --- | --- | --- | --- | --- | --- |
| CLIP (ViT-B/16) | - | 0.29 | 0.37 | 0.32 | 0.40 |
| CLIP (RN50) | - | 0.24(-0.05) | 0.30(-0.07) | 0.27(-0.05) | 0.33(-0.07) |
| WildCLIP (RN50) | B<sub>t1</sub> | 0.59(+0.30) | 0.42(+0.05) | 0.55(+0.23) | 0.38(-0.02) |
| WildCLIP | B<sub>t1</sub> | 0.60(+0.31) | 0.43(+0.06) | **0.66(+0.34)** | **0.46(+0.06)** |
| WildCLIP-LwF | B<sub>t1</sub> | 0.54(+0.25) | 0.48(+0.11) | 0.56(+0.24) | **0.46(+0.06)** |
| WildCLIP | B<sub>t1-7</sub> | 0.64(+0.35) | 0.40(+0.03) | 0.64(+0.32) | 0.37(-0.03) |
| WildCLIP-LwF | B<sub>t1-7</sub> | 0.60(+0.31) | **0.49(+0.12)** | 0.60(+0.28) | 0.44(+0.04) |
| WildCLIP-Adapter* | B<sub>t1-7</sub> → N<sub>t1</sub> | **0.67(+0.38)** | 0.38(+0.01) | 0.64(+0.32) | 0.38(-0.02) |
| WildCLIP-Adapter*-LwF | B<sub>t1-7</sub> → N<sub>t1</sub> | **0.67(+0.38)** | 0.40(+0.03) | 0.65(+0.33) | 0.41(+0.01) |
| WildCLIP-LwF-Adapter* | B<sub>t1-7</sub> → N<sub>t1</sub> | 0.54(+0.25) | 0.43(+0.06) | 0.55(+0.23) | 0.39(-0.01) |
| WildCLIP-LwF-Adapter*-LwF | B<sub>t1-7</sub> → N<sub>t1</sub> | 0.58(+0.29) | 0.45(+0.08) | 0.57(+0.25) | 0.42(+0.02) |

#### Learning without forgetting

In the previous section, we saw that training WildCLIP-LwF on the base vocabulary also improves its performance on the novel vocabulary (+0.12). We find that this effect is mainly due to the VR-LwF loss since WildCLIP alone does not have such an increase on the novel vocabulary (+0.03, Table 2). In that sense, the VR-LwF loss appears to be efficient at preserving the open-vocabulary capacities of CLIP, while limiting catastrophic forgetting. However, this increase in performance on the novel vocabulary set is compensated by a small drop in performance on the base vocabulary set. This is consistent with the idea that this loss term constrains the drift of the image embeddings by anchoring the latent space. **Adapter**: We found that the boost by the MLP adapter during the adaptation step is relatively limited (CLIP-Adapter∗ Table 1, WildCLIP-Adapter∗ Table 2). It even reduces the performance of WildCLIP-LwF-Adapter∗ (+0.12 *vs*. +0.06, Table 2). We speculate that this may be explained by the difficulty of the few-shot task on a dataset with noisy labels (*e.g*., woodland characteristics may not always be visible on image crops) and a sub-optimal training strategy.

#### Templates

We had created 10 different templates and wanted to check the impact of template augmentation. Surprisingly, training on a diverse set of caption templates does not improve the model performance on unseen templates compared to a model trained only on one template. Indeed, training with only template 1 achieves the best performance on test queries (constructed with out-of-domain templates) for both the base and the novel vocabulary (WildCLIP, Table 2). We speculate that either the expanded size of the image-caption pairs dataset complicates training, or the additional in-domain templates are themselves not suited to help the model to generalize to unseen ones.

#### Image sequences

In Tables 1 and 2, performance is computed considering every image as independent. However, camera trap images are generally taken from a sequence of multiple shots that share temporal information. Since all images do not carry the same level of information, aggregating the performance at the sequence level can further improve the performance. Appendix Table 3 shows the performance at the sequence level for CLIP, WildCLIP and WildCLIP-Adapter∗ when taking the maximum cosine similarity over the images of a sequence for each test query. As expected, we observe a consistent improvement of around +0.03 for all methods.

**Table 3.**
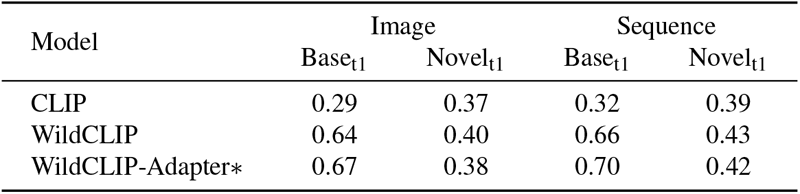
Performance at the image or sequence level.

## Discussion and Conclusion

We propose an approach based on vision-language models to retrieve scenic, behavioral, and species attributes from camera trap images with user-defined open vocabulary queries expressed as language prompts. We show that WildCLIP effectively adapts CLIP to camera traps of the Serengeti initiative and can function well to retrieve rare events of interest. We envision our method to find application in assisting the annotation process of camera trap datasets, to find rare events of interest quickly, and to facilitate species retrieval under diverse environmental conditions. This also has the potential to reduce bias when training species classifiers.

To counteract catastrophic forgetting, we adapted memory replay (20, 54) and found that it works relatively well based on a replay vocabulary mined from the scientific literature on the Serengeti. Importantly, one does not need access to the original training set or any images, which might require a lot of storage. Our results suggest that WildCLIP can retrieve events sometimes missed by CLIP for open-vocabulary queries. But the size of the Snapshot Serengeti dataset remains too limited to give any trend regarding the relative open-vocabulary performances of both models. We think this is a promising direction, and we will explore the impact of different replay vocabularies in the future. To be more reliable for the ecology community, WildCLIP would greatly benefit from a larger vocabulary and from being trained on multiple camera trap datasets. This improvement requires collaborative efforts in sharing and annotating camera trap datasets with labels that go beyond taxonomy information. We hope that our demonstration of feasibility will contribute to the emergence of more camera trap datasets that are annotated with attributes beyond species.

## Acknowledgements

We are grateful to Sepideh Mammooler for help extracting the replay vocabulary, Mackenzie Mathis for some LATEX hacks and to the ECEO and Mathis Group members for feedback as the project developed. This project was funded by EPFL’s SV-ENAC I-PhD program.

## Data Availability

Snapshot Serengeti data is publicly available on LILA BC (53). We are grateful to the authors for making this dataset public and to the actors involved in maintaining and continuously expanding LILA BC (16).

## Appendix

### Caption templates

Here we detail the full caption templates used for training WildCLIP. Templates 1 to 7 are used for training, and 1 and 8, 9, 10 for evaluation. Since we consider combinations of attributes, each template yields 31 captions.

The 10 templates:

1. A camera-trap picture of a `<age> <spe> <beh> <env> <tod>`.
2. A `<age> <spe> <beh> <env> <tod>`.
3. There is a `<age> <spe>`, it is `<beh> <env> <tod>`.
4. A picture of a `<age> <spe> <env> <tod>`, it is `<beh>`.
5. `<tod>`, a `<age>` animal is `<beh> <env>`, it is a `<spe>`.
6. An image of a `<age> <spe> <beh> <env> <tod>`.
7. `<env>`, a `<age> <spe>` is `<beh> <tod>`.
8. A wild `<spe> <beh> <env> <tod>`. It is a `<age>`.
9. In this picture, a `<age> <spe>` is `<beh> <env> <tod>`.
10. The `<age> <spe>` is `<beh> <env> <tod>`.

with:

- `<age>`: the animal age, either “young” or “adult”
- `<spe>`: the animal species name
- `<beh>`: the animal behavior
- `<env>`: the environment, either”grassland” or “woodland”
- `<tod>`: the time of the day, either “nighttime” or “daytime”

### Qualitative results for complex queries of the WildCLIP vocabulary

We tested the retrieval performance of CLIP, WildCLIP, WildCLIP-LwF and WildCLIP-LwF-adapter-LwF on multiple queries containing words from the base and novel vocabularies (Appendix Figure 10). A green box indicates that the caption describes properly the image according to the dataset labels. A red box indicates a mismatch. However, note that the labels used are noisy in nature (*cf*. the first image retrieved for the last two lines, where images depicting lions at nighttime are considered a mismatch despite being well retrieved).

**Figure 10.**
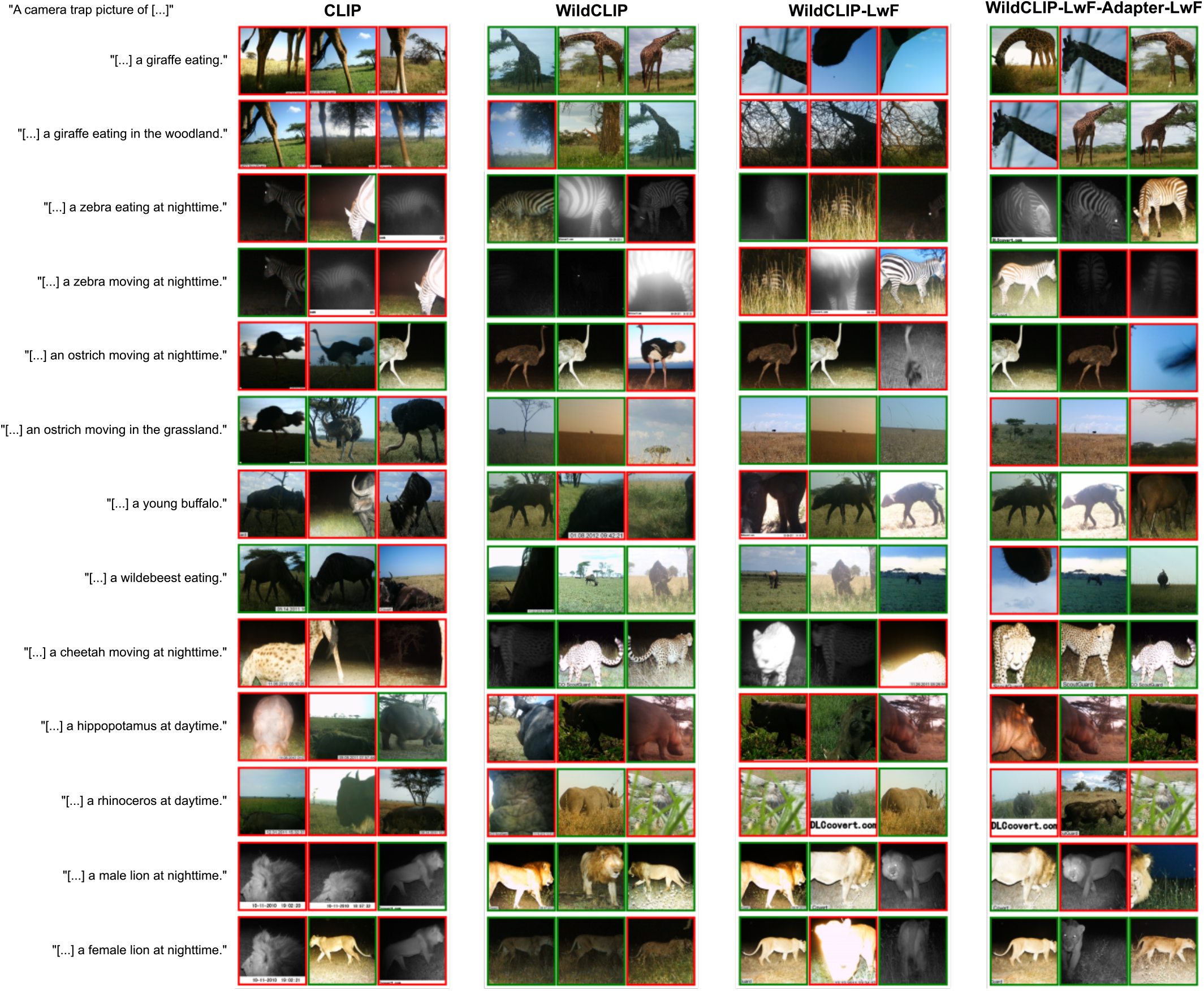
Top-3 test images most aligned with the given complex queries.

### Qualitative results for open-vocabulary queries

We tested the retrieval performance of CLIP, WildCLIP and WildCLIP-LwF on multiple queries containing words never seen during the training of WildCLIP (Appendix Figure 11). Although this ability is beyond the scope of this study, we still wish to illustrate the potential of VLM to retrieve any kind of events and to position WildCLIP with respect to this goal. We observe a decrease in performance for WildCLIP in comparison to CLIP for prompts such as *“a cloudy weather”*, but this catastrophic forgetting is compensated as expected by the VR-LwF loss. Other queries such as *“an animal eating from a tree.”* are never well retrieved, but WildCLIP seems the best model in this case since the animals are eating the closest to a tree. Finally, *“an animal with an open wound.”* is never well retrieved by any of the models, although we are aware of the presence of such images in the test set.

**Figure 11.**
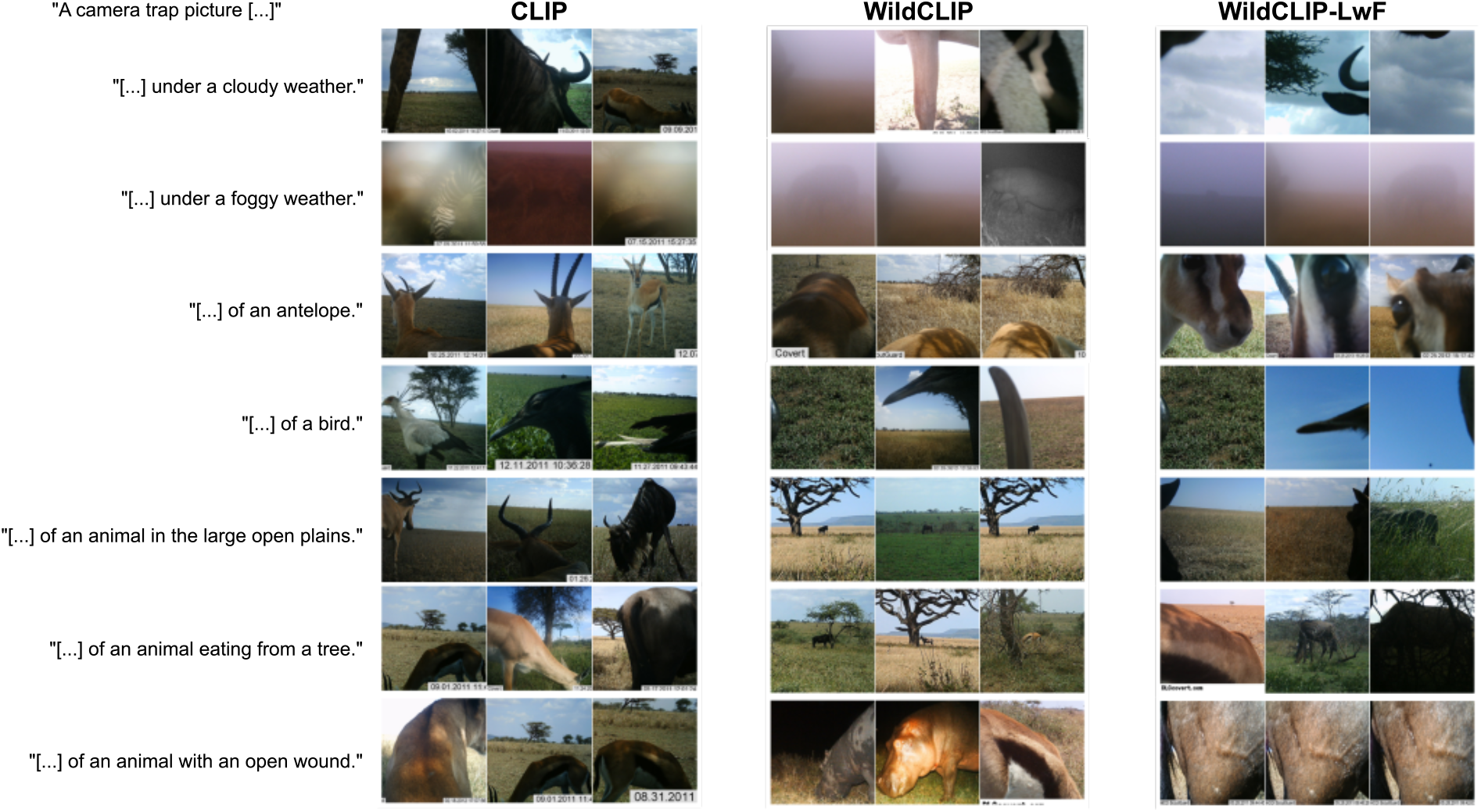
Top-3 test images most aligned with the given open-vocabulary queries.

### Sequence level performance

We compute the mAP considering either each image as independent, or by first taking the maximum cosine similarity with a given query over the sequence of camera trap shots, and then computing the mAP over the sequences. Performance is reported for the test queries of the base and the novel vocabulary set, following template 1 (Appendix Table 3).

### Implementation details

#### Data processing

We started from the output of MegaDetector (7) provided on LILA, and included bounding boxes predicted with confidence above 0.7 in our analysis. Images containing animals are then cropped to undistorted square patches by padding with background.

#### Replay vocabulary

To generate the replay-captions, we first parse titles of ecology papers corresponding to the query “Serengeti+Wildlife” with the Semantic Scholar API (55). We then use the Rapid Automatic Keyword Extraction (Rake (56)) on all titles to keep only keywords from them. Since this process is not specific to ecology, we further filter words by computing the cosine distances between words embeddings of the retrieved words and “Serengeti” and “Wildlife” using a Word2Vec model pretrained with GloVe (57). We keep only the 830 most similar words in total, and build the 5-grams text anchors by randomly sampling these words.

Examples of 5-grams used as replay vocabulary (VR):

- seasonal tasmania snakes unengaged ruminant
- coyote narok raccoons disease sustainable
- cull bird rhinoceroses act pesticide
- jaguars mammal culling territoriality canine
- feed maasai diversity poaching improve
- grass today tree browsers myxomatosis

#### Model training

During training of the different versions of WildCLIP, we use the weighted Adam optimizer (58), with a learning rate of 10^−7^ following a cosine annealing scheduler (59), a batch size of 100 and a weight decay of 0.2. The training code and parameters are adapted from (60). The learning rate is increased to 10^−6^, and the weight decay decreased to 10^−3^ for ResNet models. We randomly draw 10^*′*^000 image-caption pairs for a given epoch, with a sampling probability inversely proportional to the caption frequency. Image crops are then randomly transformed with a probability of 0.25 for horizontal flipping, resizing, Gaussian blur, conversion to grayscale and color jitter. Around 10% of the training data is used as a validation set by holding out a subset of cameras. The models are trained for 500 epochs. The training of WildCLIP-Adapter and its variants is different because of the few-shot scenario. Following parameters used in (12), we use the stochastic gradient descent (SGD) optimizer with a learning rate of 10^−3^, and train the model for 200 epochs with a batch size of 32. The *α* blending parameter in Figure 2 is set to 0.7, following cross-validated results of (12). The temperature parameter *τ* in Eq. (4, 3, 7, 7) is set to 0.01, following (60).

## References

1. Allan F O’Connell, James D Nichols, and K Ullas Karanth. Camera traps in animal ecology: methods and analyses, volume 271. Springer, 2011.

2. Robin Steenweg, Mark Hebblewhite, Roland Kays, Jorge Ahumada, Jason T Fisher, Cole Burton, Susan E Townsend, Chris Carbone, J Marcus Rowcliffe, Jesse Whittington, et al. Scaling-up camera traps: Monitoring the planet’s biodiversity with networks of remote sensors. Frontiers in Ecology and the Environment, 15(1):26–34, 2017.

3. A Cole Burton, Eric Neilson, Dario Moreira, Andrew Ladle, Robin Steenweg, Jason T Fisher, Erin Bayne, and Stan Boutin. Wildlife camera trapping: a review and recommendations for linking surveys to ecological processes. Journal of Applied Ecology, 52(3):675–685, 2015.

4. Anthony Caravaggi, Peter B Banks, A Cole Burton, Caroline MV Finlay, Peter M Haswell, Matt W Hayward, Marcus J Rowcliffe, and Mike D Wood. A review of camera trapping for conservation behaviour research. Remote Sensing in Ecology and Conservation, 3(3):109–122, 2017.

5. Devis Tuia, Benjamin Kellenberger, Sara Beery, Blair R Costelloe, Silvia Zuffi, Benjamin Risse, Alexander Mathis, Mackenzie W Mathis, Frank van Langevelde, Tilo Burghardt, et al. Perspectives in machine learning for wildlife conservation. Nature communications, 13(1):792, 2022.

6. Zackary J Delisle, Elizabeth A Flaherty, Mackenzie R Nobbe, Cole M Wzientek, and Robert K Swihart. Nextgeneration camera trapping: systematic review of historic trends suggests keys to expanded research applications in ecology and conservation. Frontiers in Ecology and Evolution, 9:617996, 2021.

7. Sara Beery, Dan Morris, and Siyu Yang. Efficient pipeline for camera trap image review. arXiv preprint 1907.06772, 2019.

8. Mohammad Sadegh Norouzzadeh, Anh Nguyen, Margaret Kosmala, Alexandra Swanson, Meredith S Palmer, Craig Packer, and Jeff Clune. Automatically identifying, counting, and describing wild animals in camera-trap images with deep learning. Proceedings of the National Academy of Sciences, 115(25):E5716–E5725, 2018.

9. Alec Radford, Jong Wook Kim, Chris Hallacy, Aditya Ramesh, Gabriel Goh, Sandhini Agarwal, Girish Sastry, Amanda Askell, Pamela Mishkin, Jack Clark, et al. Learning transferable visual models from natural language supervision. In International conference on machine learning, pages 8748–8763. PMLR, 2021.

10. Omiros Pantazis, Gabriel Brostow, Kate Jones, and Oisin Mac Aodha. Svl-adapter: Self-supervised adapter for vision-language pretrained models. arXiv preprint 2210.03794, 2022.

11. Zifeng Wang, Zhenbang Wu, Dinesh Agarwal, and Jimeng Sun. Medclip: Contrastive learning from unpaired medical images and text. arXiv preprint 2210.10163, 2022.

12. Peng Gao, Shijie Geng, Renrui Zhang, Teli Ma, Rongyao Fang, Yongfeng Zhang, Hongsheng Li, and Yu Qiao. Clip-adapter: Better vision-language models with feature adapters. arXiv preprint 2110.04544, 2021.

13. James Kirkpatrick, Razvan Pascanu, Neil Rabinowitz, Joel Veness, Guillaume Desjardins, Andrei A Rusu, Kieran Milan, John Quan, Tiago Ramalho, Agnieszka Grabska-Barwinska, et al. Overcoming catastrophic forgetting in neural networks. Proceedings of the national academy of sciences, 114(13):3521–3526, 2017.

14. Sara Beery, Grant Van Horn, and Pietro Perona. Recognition in terra incognita. In Proceedings of the European conference on computer vision (ECCV), pages 456–473, 2018.

15. Stefan Schneider, Saul Greenberg, Graham W Taylor, and Stefan C Kremer. Three critical factors affecting automated image species recognition performance for camera traps. Ecology and evolution, 10(7):3503–3517, 2020.

16. LILA BC (Labeled Image Library of Alexandria: Biology and Conservation). https://lila.science/, 2023.

17. Noa Rigoudy, Gaspard Dussert, Abdelbaki Benyoub, Aurelien Besnard, Carole Birck, Jerome Boyer, Yoann Bollet, Yoann Bunz, Gerard Caussimont, Elias Chetouane, et al. The deepfaune initiative: a collaborative effort towards the automatic identification of the french fauna in camera-trap images. bioRxiv, pages 2022–03, 2022.

18. Dan Liu, Jin Hou, Shaoli Huang, Jing Liu, Yuxin He, Bochuan Zheng, Jifeng Ning, and Jingdong Zhang. Loteanimal: A long time-span dataset for endangered animal behavior understanding. In Proceedings of the IEEE/CVF International Conference on Computer Vision, pages 20064– 20075, 2023.

19. Alexandra Swanson, Margaret Kosmala, Chris Lintott, Robert Simpson, Arfon Smith, and Craig Packer. Snapshot serengeti, high-frequency annotated camera trap images of 40 mammalian species in an african savanna. Scientific data, 2(1):1–14, 2015.

20. Yuxuan Ding, Lingqiao Liu, Chunna Tian, Jingyuan Yang, and Haoxuan Ding. Don’t stop learning: Towards continual learning for the clip model. arXiv preprint 2207.09248, 2022.

21. Tilo Burghardt and Janko Calic. Real-time face detection and tracking of animals. In 2006 8th seminar on neural network applications in electrical engineering, pages 27–32. IEEE, 2006.

22. Agnieszka Miguel, Sara Beery, Erica Flores, Loren Klemesrud, and Rana Bayrakcismith. Finding areas of motion in camera trap images. In 2016 IEEE international conference on image processing (ICIP), pages 1334–1338. IEEE, 2016.

23. Michael J Wilber, Walter J Scheirer, Phil Leitner, Brian Heflin, James Zott, Daniel Reinke, David K Delaney, and Terrance E Boult. Animal recognition in the mojave desert: Vision tools for field biologists. In 2013 IEEE Workshop on Applications of Computer Vision (WACV), pages 206–213. IEEE, 2013.

24. Xiaoyuan Yu, Jiangping Wang, Roland Kays, Patrick A Jansen, Tianjiang Wang, and Thomas Huang. Automated identification of animal species in camera trap images. EURASIP Journal on Image and Video Processing, pages 1–10, 2013.

25. Stefan Schneider, Graham W Taylor, and Stefan Kremer. Deep learning object detection methods for ecological camera trap data. In 2018 15th Conference on computer and robot vision (CRV), pages 321–328. IEEE, 2018.

26. Praneet Singh, Stacy M Lindshield, Fengqing Zhu, and Amy R Reibman. Animal localization in camera-trap images with complex backgrounds. In 2020 IEEE southwest symposium on image analysis and interpretation (SSIAI), pages 66–69. IEEE, 2020.

27. Michael A Tabak, Mohammad S Norouzzadeh, David W Wolfson, Steven J Sweeney, Kurt C VerCauteren, Nathan P Snow, Joseph M Halseth, Paul A Di Salvo, Jesse S Lewis, Michael D White, et al. Machine learning to classify animal species in camera trap images: Applications in ecology. Methods in Ecology and Evolution, 10(4):585–590, 2019.

28. Guobin Chen, Tony X Han, Zhihai He, Roland Kays, and Tavis Forrester. Deep convolutional neural network based species recognition for wild animal monitoring. In 2014 IEEE international conference on image processing (ICIP), pages 858–862. IEEE, 2014.

29. Robin C Whytock, Jedrzej ś wieżewski, Joeri A Zwerts, Tadeusz Bara-Słupski, Aurélie Flore Koumba Pambo, Marek Rogala, Laila Bahaa-el din, Kelly Boekee, Stephanie Brittain, Anabelle W Cardoso, et al. Robust ecological analysis of camera trap data labelled by a machine learning model. Methods in Ecology and Evolution, 12(6):1080–1092, 2021.

30. Marco Willi, Ross T Pitman, Anabelle W Cardoso, Christina Locke, Alexandra Swanson, Amy Boyer, Marten Veldthuis, and Lucy Fortson. Identifying animal species in camera trap images using deep learning and citizen science. Methods in Ecology and Evolution, 10(1):80–91, 2019.

31. Otto Brookes, Majid Mirmehdi, Hjalmar Kühl, and Tilo Burghardt. Triple-stream deep metric learning of great ape behavioural actions. arXiv preprint 2301.02642, 2023.

32. Michael A Tabak, Daniel Falbel, Tess Hamzeh, Ryan K Brook, John A Goolsby, Lisa D Zoromski, Raoul K Boughton, Nathan P Snow, Kurt C VerCauteren, and Ryan S Miller. Cameratrapdetector: Automatically detect, classify, and count animals in camera trap images using artificial intelligence. bioRxiv, pages 2022–02, 2022.

33. Emma R Bush, Robin C Whytock, Laila Bahaa-El-Din, Stéphanie Bourgeois, Nils Bunnefeld, Anabelle W Cardoso, Jean Thoussaint Dikangadissi, Pacôme Dimbonda, Edmond Dimoto, Josué Edzang Ndong, et al. Long-term collapse in fruit availability threatens central african forest megafauna. Science, 370(6521):1219–1222, 2020.

34. Maureen H Murray, Mason Fidino, Elizabeth W Lehrer, Juniper L Simonis, and Seth B Magle. A multi-state occupancy model to non-invasively monitor visible signs of wildlife health with camera traps that accounts for image quality. Journal of Animal Ecology, 90(8):1973–1984, 2021.

35. Craig D Reddell, Fitsum Abadi, David K Delaney, James W Cain, and Gary W Roemer. Urbanization’s influence on the distribution of mange in a carnivore revealed with multistate occupancy models. Oecologia, 195:105–116, 2021.

36. Mohammad Sadegh Norouzzadeh, Dan Morris, Sara Beery, Neel Joshi, Nebojsa Jojic, and Jeff Clune. A deep active learning system for species identification and counting in camera trap images. Methods in ecology and evolution, 12 (1):150–161, 2021.

37. Benjamin Kellenberger, Devis Tuia, and Dan Morris. Aide: Accelerating image-based ecological surveys with interactive machine learning. Methods in Ecology and Evolution, 11(12):1716–1727, 2020.

38. Benjamin Kellenberger, Diego Marcos, and Devis Tuia. Detecting mammals in uav images: Best practices to address a substantially imbalanced dataset with deep learning. Remote sensing of environment, 216:139–153, 2018.

39. Tanmay Nath, Alexander Mathis, An Chi Chen, Amir Patel, Matthias Bethge, and Mackenzie Weygandt Mathis. Using deeplabcut for 3d markerless pose estimation across species and behaviors. Nature protocols, 14(7):2152–2176, 2019.

40. Omiros Pantazis, Gabriel J Brostow, Kate E Jones, and Oisin Mac Aodha. Focus on the positives: Self-supervised learning for biodiversity monitoring. In Proceedings of the IEEE/CVF International Conference on Computer Vision, pages 10583–10592, 2021.

41. Ashish Vaswani, Noam Shazeer, Niki Parmar, Jakob Uszkoreit, Llion Jones, Aidan N Gomez, Łukasz Kaiser, and Illia Polosukhin. Attention is all you need. Advances in neural information processing systems, 30, 2017.

42. Jacob Devlin, Ming-Wei Chang, Kenton Lee, and Kristina Toutanova. Bert: Pre-training of deep bidirectional transformers for language understanding. arXiv preprint 1810.04805, 2018.

43. Colin Raffel, Noam Shazeer, Adam Roberts, Katherine Lee, Sharan Narang, Michael Matena, Yanqi Zhou, Wei Li, and Peter J Liu. Exploring the limits of transfer learning with a unified text-to-text transformer. The Journal of Machine Learning Research, 21(1):5485–5551, 2020.

44. Tom Brown, Benjamin Mann, Nick Ryder, Melanie Subbiah, Jared D Kaplan, Prafulla Dhariwal, Arvind Neelakantan, Pranav Shyam, Girish Sastry, Amanda Askell, et al. Language models are few-shot learners. Advances in neural information processing systems, 33:1877–1901, 2020.

45. Long Ouyang, Jeffrey Wu, Xu Jiang, Diogo Almeida, Carroll Wainwright, Pamela Mishkin, Chong Zhang, Sandhini Agarwal, Katarina Slama, Alex Ray, et al. Training language models to follow instructions with human feedback. Advances in Neural Information Processing Systems, 35: 27730–27744, 2022.

46. Dídac Surís, Sachit Menon, and Carl Vondrick. Vipergpt: Visual inference via python execution for reasoning. arXiv preprint 2303.08128, 2023.

47. Yongliang Shen, Kaitao Song, Xu Tan, Dongsheng Li, Weiming Lu, and Yueting Zhuang. Hugginggpt: Solving ai tasks with chatgpt and its friends in huggingface. arXiv preprint 2303.17580, 2023.

48. Shaokai Ye, Jessy Lauer, Mu Zhou, Alexander Mathis, and Mackenzie W Mathis. AmadeusGPT: a natural language interface for interactive animal behavioral analysis. Advances in Neural Information Processing Systems, 2023.

49. Jean-Baptiste Alayrac, Jeff Donahue, Pauline Luc, Antoine Miech, Iain Barr, Yana Hasson, Karel Lenc, Arthur Mensch, Katherine Millican, Malcolm Reynolds, et al. Flamingo: a visual language model for few-shot learning. Advances in Neural Information Processing Systems, 35:23716–23736, 2022.

50. Jiasen Lu, Dhruv Batra, Devi Parikh, and Stefan Lee. Vilbert: Pretraining task-agnostic visiolinguistic representations for vision-and-language tasks. Advances in neural information processing systems, 32, 2019.

51. Chao Jia, Yinfei Yang, Ye Xia, Yi-Ting Chen, Zarana Parekh, Hieu Pham, Quoc Le, Yun-Hsuan Sung, Zhen Li, and Tom Duerig. Scaling up visual and vision-language representation learning with noisy text supervision. In International conference on machine learning, pages 4904–4916. PMLR, 2021.

52. Zhizhong Li and Derek Hoiem. Learning without forgetting. IEEE transactions on pattern analysis and machine intelligence, 40(12):2935–2947, 2017.

53. Snapshot Serengeti labeled information, library of alexandria: Biology and conservation website. https://lila.science/datasets/snapshot-serengeti, 2019.

54. Shaokai Ye, Anastasiia Filippova, Jessy Lauer, Maxime Vidal, Steffen Schneider, Tian Qiu, Alexander Mathis, and Mackenzie Weygandt Mathis. Superanimal models pretrained for plug-and-play analysis of animal behavior. arXiv preprint 2203.07436, 2022.

55. Rodney Kinney, Chloe Anastasiades, Russell Authur, Iz Beltagy, Jonathan Bragg, Alexandra Buraczynski, Isabel Cachola, Stefan Candra, Yoganand Chandrasekhar, Arman Cohan, et al. The semantic scholar open data platform. arXiv preprint 2301.10140, 2023.

56. Stuart Rose, Dave Engel, Nick Cramer, and Wendy Cowley. Automatic keyword extraction from individual documents. Text mining: applications and theory, pages 1–20, 2010.

57. Jeffrey Pennington, Richard Socher, and Christopher D Manning. Glove: Global vectors for word representation. In Proceedings of the 2014 conference on empirical methods in natural language processing (EMNLP), pages 1532– 1543, 2014.

58. Ilya Loshchilov and Frank Hutter. Decoupled weight decay regularization. arXiv preprint 1711.05101, 2017.

59. Ilya Loshchilov and Frank Hutter. Sgdr: Stochastic gradient descent with warm restarts. arXiv preprint 1608.03983, 2016.

60. Gabriel Ilharco, Mitchell Wortsman, Ross Wightman, Cade Gordon, Nicholas Carlini, Rohan Taori, Achal Dave, Vaishaal Shankar, Hongseok Namkoong, John Miller, Hannaneh Hajishirzi, Ali Farhadi, and Ludwig Schmidt. Openclip, July 2021.

